# Intercellular collectivity is governed by enzyme secretion strategies in marine polysaccharide degrading bacteria

**DOI:** 10.1101/2022.04.19.488775

**Authors:** Glen D’Souza, Ali Ebrahimi, Astrid Stubbusch, Michael Daniels, Johannes Keegstra, Roman Stocker, Otto Cordero, Martin Ackermann

**Affiliations:** Microbial Systems Ecology Group, Institute of Biogeochemistry and Pollutant Dynamics, Department of Environmental Systems Sciences, ETH Zurich, 8006 Zurich, Switzerland; Department of Environmental Microbiology, Eawag: Swiss Federal Institute of Aquatic Sciences, 8600 Duebendorf, Switzerland; Department of Civil and Environmental Engineering, MIT, Cambridge, MA 02139, USA; Institute of Environmental Engineering, Department of Civil, Environmental and Geomatic Engineering, ETH Zurich, 8093 Zurich, Switzerland

**Keywords:** marine bacteria, enzyme secretion, dispersal, density dependence, intercellular collectivity, polysaccharide degradation

## Abstract

Polysaccharide breakdown by bacteria requires the activity of enzymes that degrade polymers extracellularly. This generates a localized pool of breakdown products that are accessible to the enzyme producers themselves as well as to other organisms. Marine bacterial taxa often show marked differences in the production and secretion of degradative enzymes that break down polysaccharides. These differences can have profound effects on the pool of diffusible breakdown products and hence on the ecological dynamics. However, the consequences of differences in enzymatic secretions on cellular growth dynamics and interactions are unclear. Here we combine experiments and models to study the growth dynamics of single cells within populations of marine *Vibrionaceae* strains that grow on the abundant marine polymer alginate, using microfluidics coupled to quantitative single-cell analysis and mathematical modelling. We find that strains that have low extracellular secretions of alginate lyases show stronger aggregative behaviors compared to strains that secrete high levels of enzymes. One plausible reason for this observation is that low secretors require a higher cellular density to achieve maximal growth rates in comparison with high secretors. Our findings indicate that increased aggregation increases intercellular synergy amongst cells of low-secreting strains. By mathematically modelling the impact of the level of degradative enzyme secretion on the rate of oligomer loss to diffusion, we find that enzymatic capability modulates the propensity of cells within clonal populations to cooperate or compete with each other. Our experiments and models demonstrate that marine bacteria display distinct aggregative behaviors and intercellular interactions based on their enzymatic secretion capabilities when growing on polysaccharides.

## Introduction

Complex polysaccharides represent the largest pools of metabolizable resources that microbial populations encounter in their environment. The degradation of these polysaccharides by bacteria drives major biogeochemical processes such as the remineralization of particulate organic matter^1–3^. Polysaccharides have high molecular weights and are thus degraded by enzymes that act extracellularly, thereby generating a pool of diffusible metabolites^1,3^. As a result, the degradative activity of one cell has the potential to influence the growth and metabolism of neighboring cells^1,4–6^. However, the byproducts generated due to extracellular digestion can be lost through diffusion^7–9^. In order to reduce diffusional losses, cells can aggregate and thereby increase local cell densities^10–12^. This group behavior can allow cells to benefit from the degradative activities of other cells and enhance growth of the clonal population while growing on polysaccharides^11^.

Closely related strains can show marked differences in the secretion of enzymes that are required to degrade polysaccharides^13^. For instance, marine *Vibrionaceae* strains differ in the secretion of extracellular alginate lyases that degrade alginate, a marine algal-derived polysaccharide composed of alternating units of guluronic acid and mannuronic acid^13,14^. While it is known that the level of secretion of alginate lyases by *Vibrionaceae* strains influences population growth dynamics in well-mixed environments^13^, it is generally unclear how differences in enzymatic secretion influence the behavior of individual cells. Studying these dynamics at the microscale will provide a better understanding of intercellular interactions within bacterial aggregates, a growth behavior commonly observed when bacteria grow on polysaccharides^10,11,14,15^. In this study, we sought to address these knowledge gaps by asking how the level of alginate lyase secretion influences aggregation and the growth dynamics of cells growing on polysaccharides.

We studied 12 marine *Vibrionaceae* isolates that have previously been shown to differ in the types and number of alginate lyases encoded in their genome^13^. They also differ in the total amount of these enzymes that are produced and secreted^13,14^. These strains were previously isolated from marine habitats and span three distinct species: *Vibrio splendidus, Vibrio cyclitrophicus* and *Vibrio sp. F13^13^* (Supplementary Table 1). We first studied well-mixed batch cultures to determine whether population growth dynamics of the *Vibrionaceae* isolates are associated with the amount of alginate lyases that are produced and secreted. We then used microfluidics coupled to time-lapse microscopy^11^ in order to quantify aggregation and single-cell growth dynamics of strains growing on alginate. Finally, we used mathematical modelling to ascertain the impact of enzyme activity on polysaccharide breakdown and cellular interactions. Together, these experiments and models show that differences in the secretion of alginate lyases between strains are associated with differences in the aggregative behaviors and density-dependence of growth rates of bacterial populations.

## Results and Discussion

### Strains belonging to the same species display distinct growth dynamics on the marine polysaccharide alginate

We first quantified the growth dynamics of the 12 *Vibrionaceae* populations on alginate in well-mixed batch cultures. Growth of populations was initiated at approximately the same inoculum density (10^5^ colony forming units (c.f.u.) ml^-1^). We tracked the growth dynamics by measuring the optical density at 600 nm and compared the maximum population size reached over the course of 36 h (Figures 1 and S1). We found significant differences in the maximal optical density achieved by different strains within each species (Figures 1 and S1). In *V. splendidus,* strains 12B01 and FF6 reached a lower maximum population size compared to strains 1S124 and 13B01 (Figures 1 and S1A). In *V. cyclitrophicus,* strain ZF270 reached a lower maximum population size compared to strains 1F175, 1F111, and ZF28 (Figures 1 and S1A). Similarly, in *V. sp.* F13, strain 9ZC77 reached a lower maximum population size than strains 9CS106, 9ZC13, and ZF57 (Figures 1 and S1A). These findings suggest that some strains are limited in their growth abilities in well-mixed environments, perhaps as a consequence of differences in the amount and activity of enzymes they release (Supplementary Table 1).

**Figure 1:**
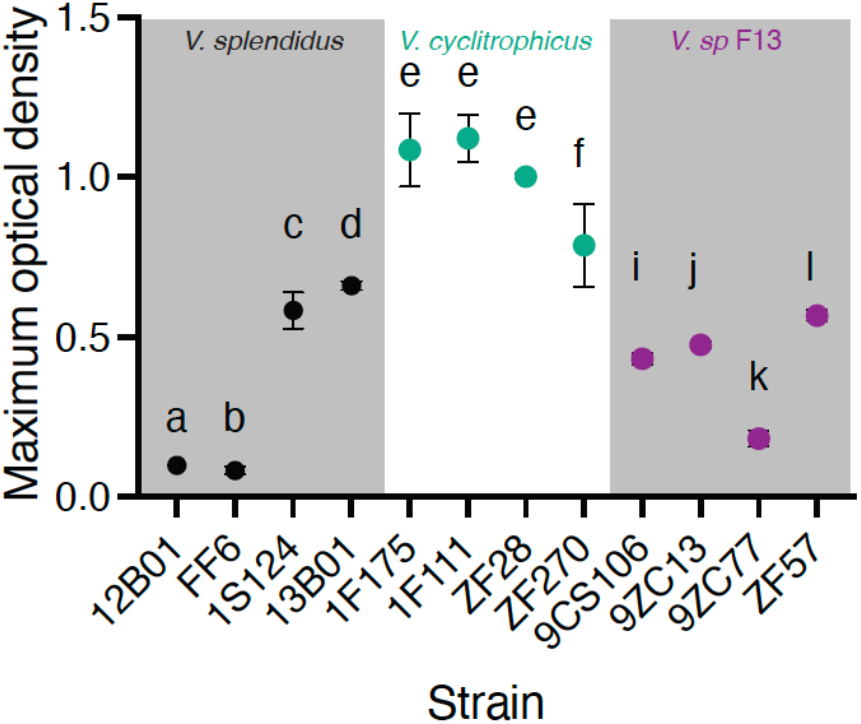
12 *Vibrionaceae* strains differ in their maximum growth on the marine polysaccharide alginate under well-mixed conditions. Maximum optical density (measured at 600 nm) achieved by strains belonging to *Vibrio splendidus, Vibrio cyclitrophicus* and *Vibrio sp.* F13 during the course of a 36 h growth cycle on the same concentration (0.1 % weight/volume) of the polysaccharide alginate. Circles and error bars indicate the mean of measurements across populations within each strain (*n*_populations_ = 3) and the 95% confidence interval (CI), respectively. Different letters indicate statistically significant differences between strains within the same species (one-way ANOVA and Dunnett’s post-hoc test; *V. splendidus: P* < 0.0001, *F* = 325.9, R^2^ = 0.99; *V. cyclitrophicus: P* = 0.0099, *F* = 7.60, R^2^ = 0.74; *V. sp* F13: *P* < 0.0001, *F* = 245, R^2^ = 0.98). For growth curves of individual strains, see Figure S1.

### Differences in enzyme secretion are associated with differences in the growth dynamics of alginate degraders

We hypothesized that reduced secretion of alginate lyases is responsible for limiting the growth of strains that achieve low optical densities in batch culture (Figure 1). To test our prediction, we measured the level of secretions of alginate degrading enzymes by the different strains using a plate assay. In this assay, alginate reacts with iodine to produce a violet compound. The breakdown of alginate creates simpler oligomers or monomers that yield colorless halos when stained with iodine (Figure S2A). Therefore, the level of enzyme secretion and activity can be quantified by measuring the size of the alginate-free halo generated by cell colonies as they grow on an alginate-containing plate (see Methods and Figure S2A)^16^.

Enzyme production as measured by halo diameter varied among strains within each species. Among *V. splendidus* strains, 12B01 and FF6 had the lowest halo diameters (Figure S2B), while among *V. cyclitrophicus* strains, ZF28 had a smaller halo diameter than the other strains (Figure S2B). Interestingly, in contrast to the other *V. sp.* F13 strains, 9ZC77 did not produce a detectable halo (Figure S2B). This observation suggests that 9ZC77 either does not produce substantial enzymatic secretions or the alginate lyases are primarily membrane-bound. Comparison of the results from the plate assays (Figure S2) and the batch cultures (Figure 1) reveals that the level of secretion of alginate lyases by the different strains is significantly correlated with the maximum population size they achieved when growing on alginate under well-mixed conditions (Figure 2A). This correlation supports our hypothesis that growth in batch culture is limited by the ability to produce alginate lyases.

**Figure 2:**
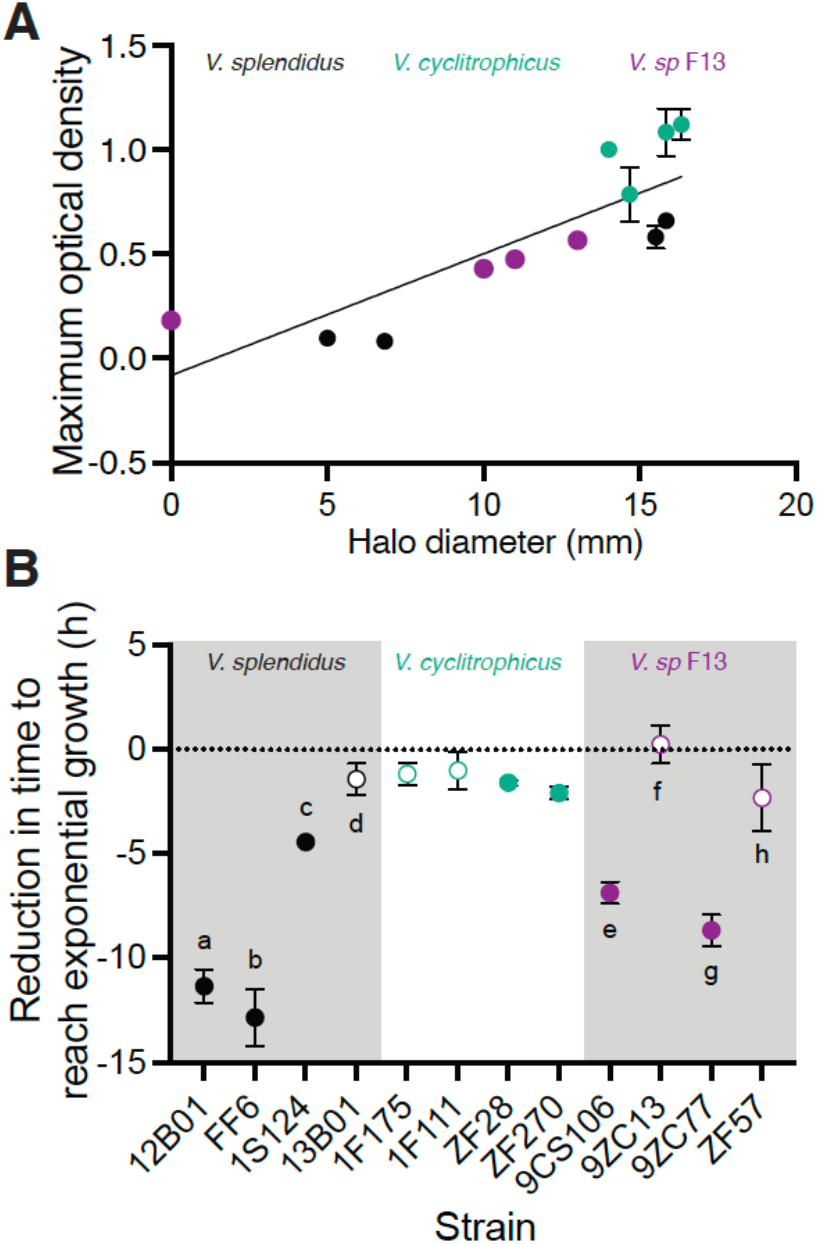
Enzyme secretion capabilities govern growth dynamics of alginate degraders. (**A**) Correlation between maximum optical density in batch culture (Figure 1) and halo diameters in the plate assays across the 12 strains. The line indicates the fit of a linear regression (slope = 0.05, R^2^ = 0.67, *P* < 0.0001; Spearman’s correlation: *r* = 0.90, *P* = 0.0002). **(B)** Supplementation with external alginate lyase reduces the time to reach the exponential growth phase of strains that secrete low levels of alginate lyase to a greater extent compared to high secretors of alginate lyase. Circles represent the mean of the populations of each ecotype (*n*_populations_ = 3) and error bars indicate 95% confidence intervals. Letters indicate statistically significant differences between strains within each species (Oneway ANOVA with Dunnett’s posthoc test; *V. splendidus: F* = 114.0, *P* < 0.0001, *R^2^* = 0.97; *V. cyclitrophicus: F* = 2.36, *P* = 0.14, *R^2^* = 0.46; *V. sp.* F13: *F* = 47.27, *P* < 0.0001, *R^2^* = 0.94). Open circles indicate non-significant differences from 0, indicating no reduction in the time to reach exponential phase; filled circles indicate significant differences from 0 (One sample *t*-test; 12B01: *t =* 24.43, df = 2, *P* = 0.001; FF6: *t* = 16.14, df = 2, *P* = 0.003; 1S124: *t* = 53, df = 2, *P* = 0.004; 13B01: *t* = 3.21, df = 2, *P* = 0.08; 1F175: *t* = 3.88, df = 2, *P* = 0.06; 1F111: *t* = 1.92, df = 2, *P* = 0.19; ZF28: *t* = 19, df = 2, *P* = 0.002; ZF270: *t* = 3.88, df = 2, *P* = 0.006; 9CS106: *t* = 23.04, df = 2, *P* = 0.0019; 9ZC13: *t* = 0.48, df = 2, *P* = 0.67; 9ZC77: *t* = 19.65, df = 2, *P* = 0.002; ZF57: *t* = 2.47, df = 2, *P* = 0.13). See Figure S3 for growth dynamics of each ecotype on alginate as well as correlations between enzyme secretion and growth kinetics.

To further test our prediction that alginate lyase production capability constrains growth, we determined whether externally supplementing alginate lyases could alleviate the growth limitation of the strains that are low enzyme secretors. We grew the 12 strains individually as batch-cultures in alginate medium that was supplemented with a commercial mixture of alginate lyases and measured their growth dynamics. The results show that alginate lyase supplementation did not alter the maximum population size reached by the low secreting strains (Figure S3A). We then calculated the time to reach exponential growth phase of each ecotype when growing in the presence of external alginate lyases and compared this with the time taken in the absence of this external supplementation. We found that the strains with a low level of enzyme secretion and activity, such as 12B01, FF6 and 9ZC77, showed the largest reduction in the time required to reach exponential phase upon supplementation with external alginate lyases (Figure 2B and S3BC). The impact of enzyme supplementation was more limited in *V. cyclitrophicus* (Figure 2A), the species in which differences in enzyme activity were most limited (Figure S2B, S3A and D). Our results thus suggest that strains that secrete low levels of alginate lyases benefit the most from external alginate lyase supplementation, indicating that the growth rate of these strains is limited by the ability to produce alginate lyases.

### Strains with low levels of enzyme secretion display higher levels of aggregation in environments that favor spatial structure

Cell aggregation can reduce the growth limitations observed on polysaccharides^11^. Specifically, we expect that aggregation could reduce the diffusional loss of alginate lyases or breakdown products and thereby could alleviate growth limitation of strains with low secretion levels. We tested this prediction on a subset of the original 12 *Vibrionaceae* strains. From each species, we selected a representative ecotype producing high levels of alginate lyase *(V.splendidus,* 13B01; *V. cyclitrophicus,* 1F111; and *V. sp.* F13, 9ZC13) and one producing low levels of alginate lyase *(V.splendidus,* 12B01; *V. cyclitrophicus,* ZF270; and *V. sp.* F13, 9ZC77). To test if strains aggregate when growing on the polysaccharide alginate, we grew cells of the different strains within microfluidic growth chambers. Within these chambers cells receive a constant supply of alginate and divide, either forming aggregates by growing as monolayers or dispersing after division. This approach enabled the quantification of growth and movement using time-lapse microscopy coupled to automated image analysis.

We found that aggregative behaviors were common in all the strains across the three species tested (Figure 3A-F). By following the growth dynamics of individual cells using cell segmentation and tracking, we were able to map lineages of all cells present in each growth chamber based on division events. This analysis revealed that cells emerging from the same founder cell tended to remain close to each other (Figure 3A-F, Supplementary Video 1-6). However, for all three species, the low-secreting strains, (Figure 3G-I), displayed a higher degree of aggregation compared to the corresponding high-secreting strains, 13B01 (Figure 3G), 1F111 (Figure 3H), and 9ZC13 (Figure 3I). The maximum number of cells within aggregates in a chamber was on average 5.54, 3.21, and 1.89 times higher for the low-secreting strains of *V. splendidus, V. cyclitrophicus,* and *V. sp* F13, respectively, compared to the corresponding high-secreting strains (Figure S4). These findings indicate that cells of low-secreting strains, at least in the context of the three species-pairs tested, form larger aggregates while growing on alginate.

**Figure 3:**
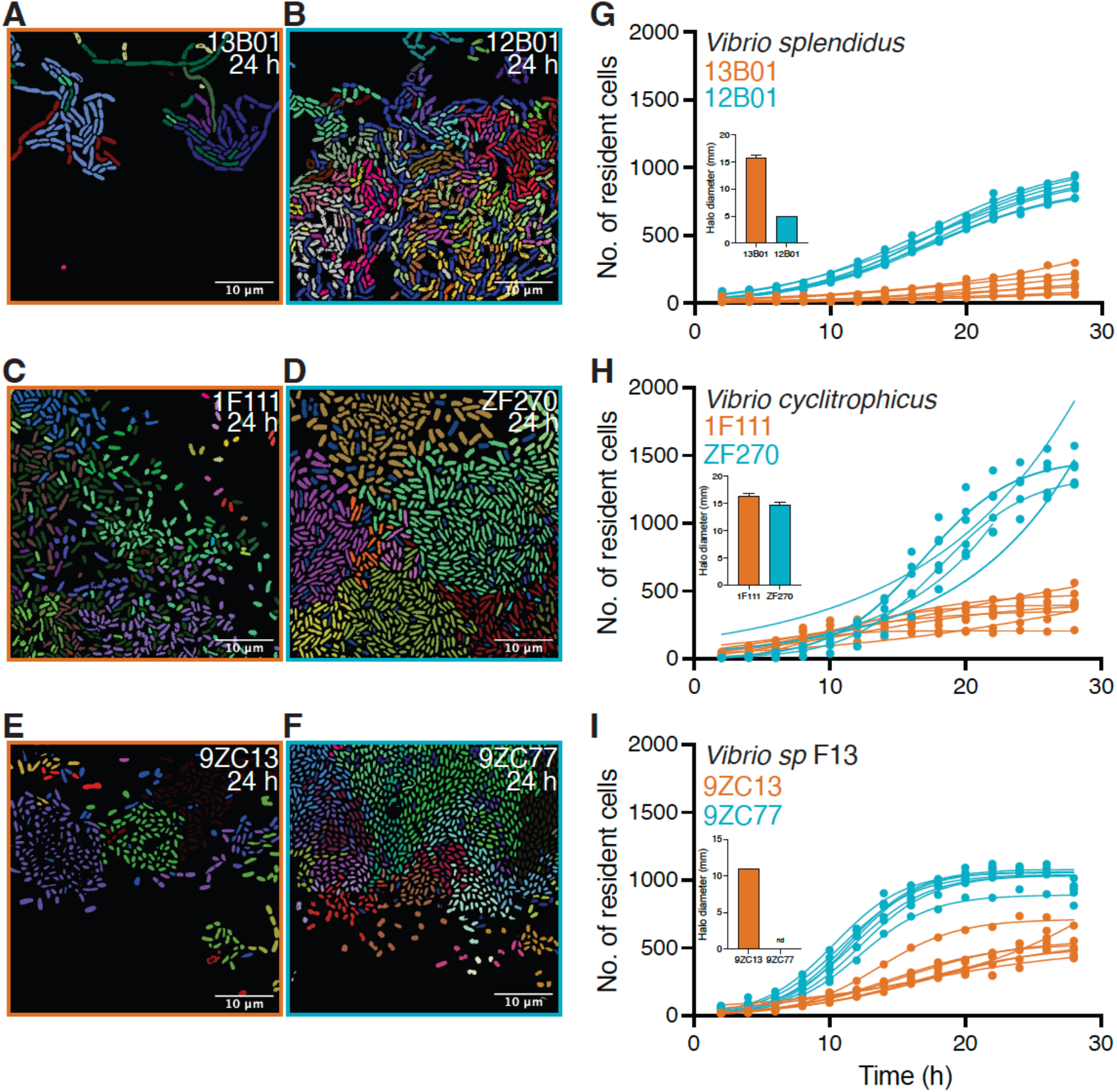
Low enzymatic production is associated with stronger aggregation. Representative images of microfluidic growth chambers containing cells of (**A**) *Vibrio splendidus* 13B01, (**B**) *Vibrio splendidus* 12B01, (**C**) *Vibrio cyclitrophicus* 1F111, (**D**) *Vibrio cyclitrophicus* ZF270, (**E**) *Vibrio sp.* F13 9ZC13, and (**F**) *Vibrio sp.* F13 9ZC77. Cells that originate from the same founder cell are shown in the same color (based on cell tracking through divisions). The three high-secreting strains (13B01, 1F111 and 9ZC13) are shown with orange panel outlines and the three high-alginate producing strains (12B01, ZF270 and 9ZC77) are shown with blue panel outlines. (**G–I**) The increase in the number of resident cells in growth chambers over time for strains of (**G**) *V. splendidus,* (**H**) *V. cyclitrophicus* and (**I**) *Vibrio sp.* F13. The levels of enzymatic secretions for the strains (from Figure S2B) are shown in the inset of panels **G-I**. The number of cells in a chamber depends on both the rate at which cells divide and the fraction of cells that leaves the chamber. Circles indicate the number of cells present at a given time point in each chamber (*n*_chambers_ = 7), with a logistic growth regression line for each chamber: 13B01, *R*^2^ = 0.97–0.99, max cells (mean ± standard deviation) = 156.90 ± 83.21, *k* = 0.09–0.20 h^-1^; 12B01, *R^2^* = 0.99 in all cases, max cells = 858.70 ± 67.78, *k* = 0.18–0.20 h^-1^; 1F111, *R*^2^ = 0.89–0.99, max cells = 410.40 ± 106.40, *k* = 0.10–0.53 h^-1^; ZF270, *R*^2^ = 0.86–0.99, max cells = 1318 ± 198.70,*k* = 0.10–0.30 h^-1^; 9ZC13, *R*^2^ = 0.91–0.99, max cells = 545.40 ± 50.69, *k* = 0.08–0.21 h^-1^; 9ZC77, *R*^2^ = 0.98–0.99, max cells = 1033 ± 72.12, *k* = 0.36–0.40 h^-1^). See supplemental videos S1-S6 for timelapse image series of cells.

### Density-dependence is stronger in strains with low alginate lyase secretions

Given that low secretion by strains is associated with greater aggregation, we then quantified if the differential aggregation behaviors of strains influence growth abilities of individual cells on the polysaccharide alginate. For this, we measured the growth rate of individual cells within microfluidic chambers using automated image analysis (Figure S5-C). We then analyzed the relationship between the median growth rate of all the cells present in a chamber during a given time interval and the total number of cells that were present in the chamber during that interval. For cells of all six strains across three species, the median growth rate increases with the increase in the number of resident cells in a chamber up to a point, and then decreases as the number of cells increases further (Figure 4). However, the number of cells required to achieve the maximal growth rate differed between the strains. For two of the three species tested, the maximal growth rate of the low-secreting ecotype occurred at a significantly higher cell density than that of the high-secreting ecotype (Figure 4A,B and E,F). The comparison for *V. cyclitrophicus* did not follow this trend, with no significant difference between the two *V. cyclitrophicus* strains in the cell density associated with the maximal growth rate (Figure 4C,D), likely because the two strains are much more similar in their level of secretion in comparison with the other species (Figure 2A). The increased cell density due to aggregation within the microfluidic chambers allows cells to overcome growth limitations that strains face in well-mixed environments (Figure S6A). Well-mixed environments, as in the batch-culture experiments, limit the growth rate of the two strains with the lowest levels of enzyme secretion, 12B01 and 9ZC77, relative to their high-secreting counterparts, 13B01 and 9ZC13, respectively (Figure S6B). In contrast, when growing in spatially structured environments, as in the microfluidic growth chambers, 12B01 and 9ZC77 are able to improve growth substantially relative to 13B01 and 9ZC13, respectively (Figure S6B). In contrast, the growth rate of *V. cyclitrophicus* ecotype ZF270 relative to that of ecotype 1F111 is similar in both well-mixed and microfluidic growth environments (Figure S6B). These findings indicate that aggregation allows strains, especially those that secrete very low levels of alginate lyases, to increase cell density and thereby enable intercellular synergy amongst cells within aggregates. The increased cell density thus enables cells in spatially structured environments to overcome growth limitations that arise when growing on polysaccharides in well-mixed environments.

**Figure 4.**
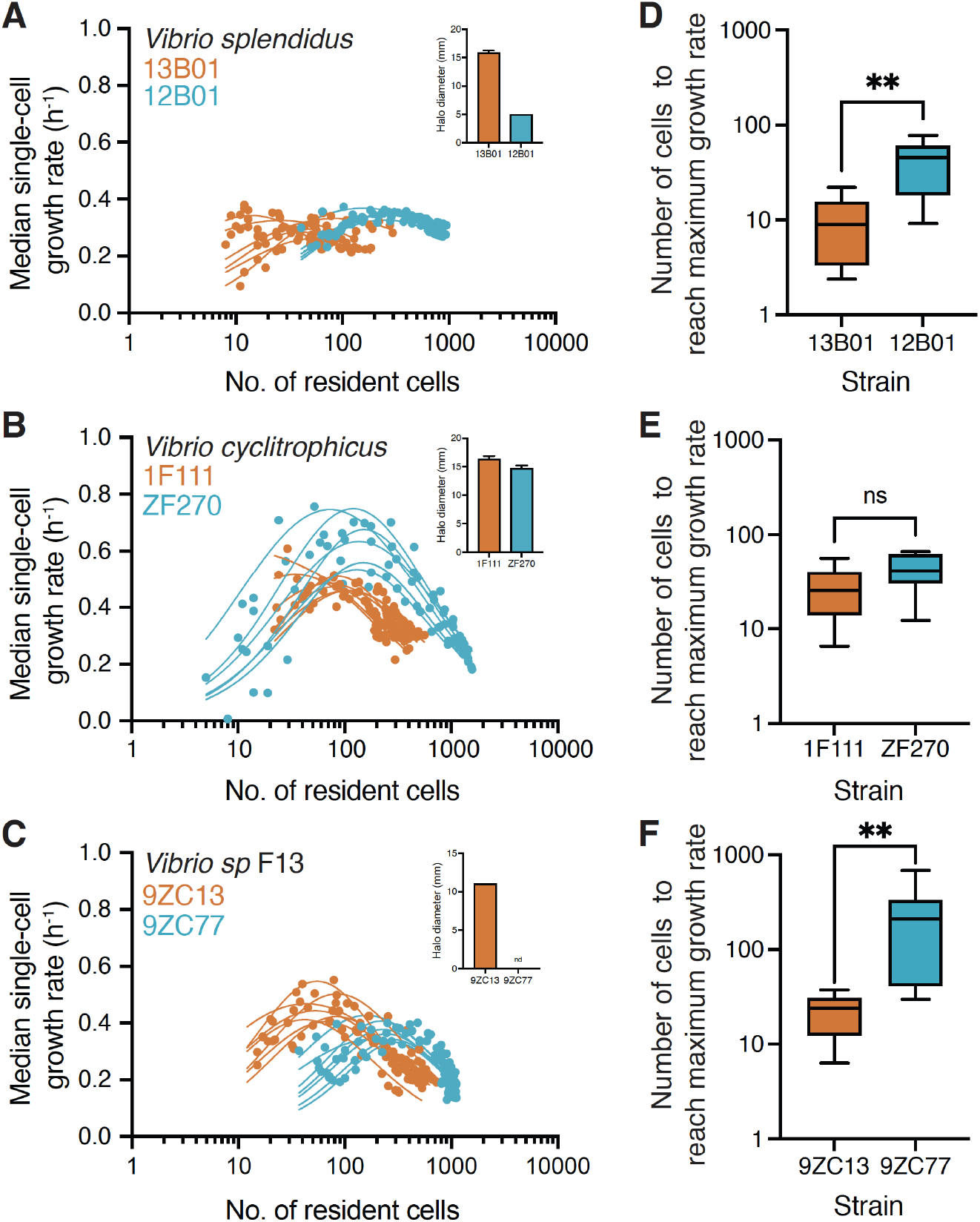
Strains with low levels of enzyme secretion rely on an increased intercellular synergy during aggregative growth. (**A, B, C**) Median single-cell growth rate (h^-1^) of (**A**) *Vibrio splendidus,* (**B**) *Vibrio cyclitrophicus* and (**D**) *Vibrio sp.* F13 cells as a function of the number of cells growing together in a microfluidic chamber. Cells were binned into 2-hour intervals based on their birth times over a 28 h experiment. This approach also yielded the number of cells present during growth in each time interval. The relationship between median growth rate of all cells present in a 2 h interval in a given chamber and the number of cells in the chamber was determined using a non-linear regression model for each chamber. The levels of enzymatic secretions for the strains (from Figure S2B) are shown in the inset of panels. High-secretion strains are indicated in orange whereas relatively low-secretion strains are indicated in blue. Circles represent data for a single bin from one chamber, and lines indicate the trajectory of growth rates for each chamber (*n*_chambers_ = 7; 13B01, *R*^2^ = 0.52–0.76; 12B01, *R*^2^ = 0.66– 0.95; 1F11, *R*^2^ = 0.88–0.91; ZF270, *R*^2^ = 0.71–0.93; 9ZC13, *R*^2^ = 0.73–0.95; 9ZC77, *R*^2^ = 0.57–0.80). (**D, E, F**) Number of cells required to reach half of the maximum growth rate for strains of (**D**) *V. splendidus,* (**E**) *V. cyclitropicus,* and (**F**) *V. sp* F13. The low-secreting *V. splendidus* ecotype 12B01 required a five-fold higher number of cells to achieve maximal growth rates compared to the high-secreting 13B01 cells (**D**; 13B01 median 8.99 cells and 12B01 median 45.25 cells). In the case of *V. cyclitrophicus,* both strains required a similar number of cells to reach maximal growth rates (**E**; 1F111 median 25.48 cells and ZF270 median 41.34 cells). For *V. sp.* F13, the low-secreting ecotype 9ZC77 required an eight-fold higher cell number to achieve maximal growth rates compared to the high-secreting ecotype 9ZC13 (**F**; 9ZC13 median 23.90 cells and 9ZC77 median 210.40 cells). Asterisks indicate statistically significant differences (*P* < 0.001) between strains (Mann–Whitney tests, *n*_chambers_ = 7; *V. splendidus, P* = 0.008, Hodges–Lehmann difference = 36.75; *V. cyclitrophicus P* = 0.15, Hodges–Lehmann difference = 19.85; *V. sp* F13 *P* = 0.003, Hodges–Lehmann difference = 188.5). Box plots extend from the 25th to 75th percentiles and whiskers indicate the 10th and 90th percentiles of median growth rates.

Once the growth rate has reached its maximum value, the growth rate then plateaus for *V. splendidus* strains or begins to decrease for *V. cyclitrophicus* and *V. sp* F13 strains with further increase in the cell numbers within the growth chambers (Figure 4A, C and E). We measured the peak cell densities reached within the growth chambers (13B01: 250.30 ± 129 (mean cell number across replicate chambers ± standard deviation), 12B01: 1439 ± 98.95, 1F111: 255 ± 42.85, ZF270: 418.60 ± 34.11, 9ZC13: 228.80 ± 39.61 and 9ZC77: 358.5 ± 109.10) and found that the benefit of intercellular interactions is balanced (for *V. splendidus*) or outweighed (for *V. cyclitrophicus* and *V. sp.* F13) by the negative effects of competitive interactions between cells within aggregates.

We then probed if an increased initial inoculum density would allow low-secreting strains to increase growth rates in well-mixed batch culture environments. In line with our findings in microfluidic chambers, low-secreting strains 12B01 and 9ZC77 benefit from a higher inoculum density (Figure S7A-C). In contrast, an increased inoculum density reduces the growth rate of high-secreting strains in well-mixed environments (Figure S7A-C). These findings lend further credence to our idea that high cell densities due to aggregation can benefit low enzyme secreting strains.

### A mathematical model offers a mechanistic explanation for the role of cell density and enzyme secretion on polymer degradation and growth rates

To obtain a mechanistic understanding of the roles of cell density and enzyme secretion rate on growth rates of strains, we developed an individual-based model of the growth rate of cells on polysaccharides taking into account cell density, enzymatic breakdown and loss of breakdown products. Our goal was to describe a system composed of spatially structured populations in nature similar to cells growing as a monolayer in microfluidic growth chambers. We modeled a 2D system (Supplementary Information) where cells occupy sites on a 50 × 50 grid (Figure 5A). We assumed diffusion as the mode of transport within the 2D chamber for polymeric substrates, secreted enzymes, and breakdown products. The loss of breakdown products (oligomers) and enzymes from the opening of the chamber by flow is simulated by assuming constant zero concentrations as the boundary condition. Individual cells are initially uniformly distributed on the 2D lattice, where they secrete enzymes that breakdown polysaccharides, consume oligomers, and grow in response to local oligomer concentrations. Monod-type kinetics were used to model substrate uptake and cell growth (see methods for details). The rate of enzyme secretion, *S*_E_, was assumed to follow first-order kinetics as a function of cell biomass, *S_E_* = *K_enz_B*, where *K_enz_* is the enzymatic activity and an intrinsic cellular property and *B* is the individual cell biomass.

**Figure 5.**
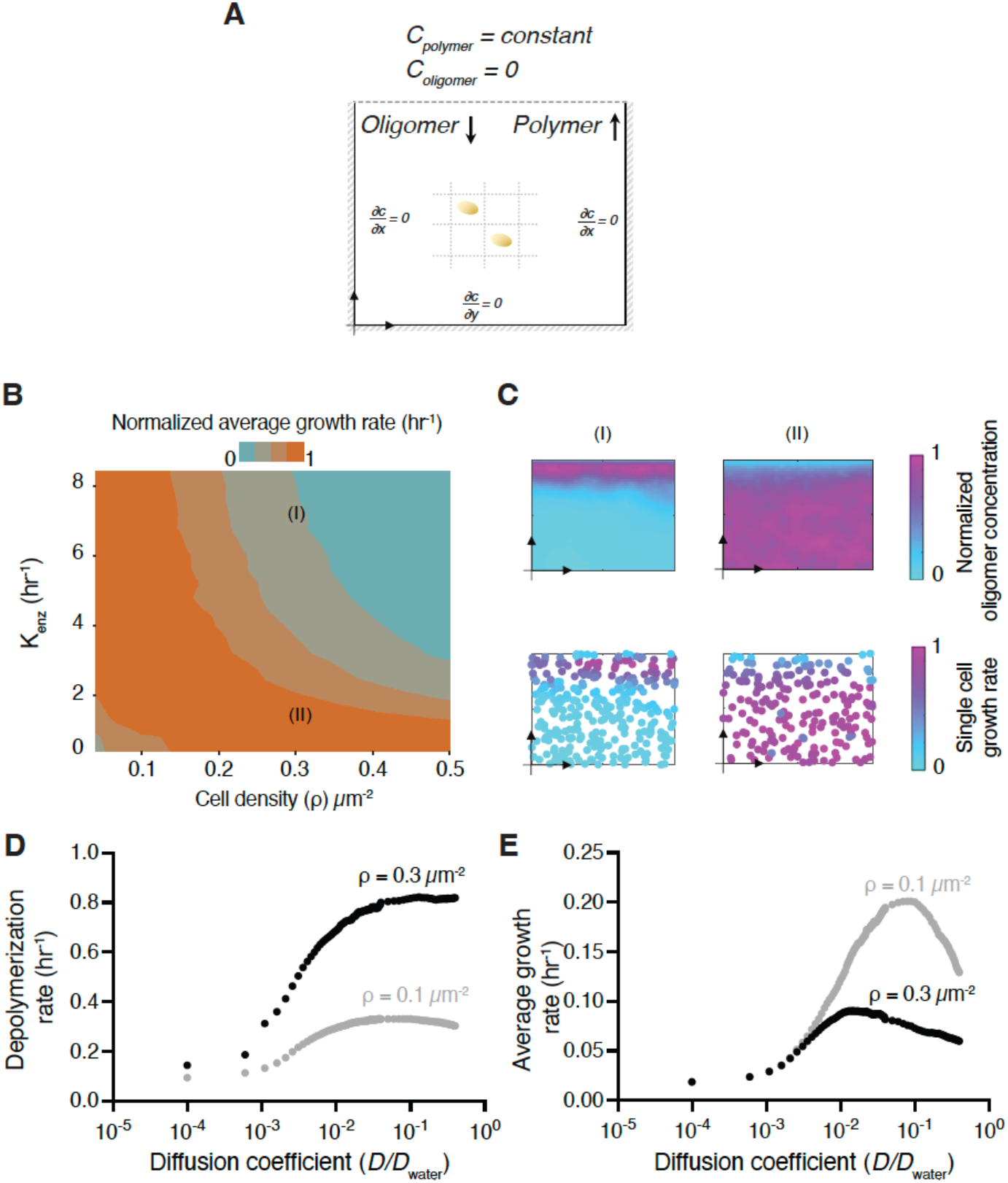
A mathematical model illustrates the biophysical mechanisms underlying bacterial density-dependent growth on polysaccharides. (A) Schematic representation of the mathematical model that simulates microfluidic chambers as a 2D lattice with 50 × 50 μm grid. Individual cells occupy single grid cells where they act as sink terms for the oligomer diffusion-reaction model. A Dirichlet boundary condition is used at the chamber periphery to simulate the effect of flow, which allows for constant concentrations of polysaccharide (0.1 mg/L) and oligomer (0 mg/L) at the boundary. Cells are uniformly distributed in the chamber. (B) Average growth rates of individual cells 20 h after the start of the simulations as a function of enzymatic activity, *K*_enz_, and cell density, *ρ*. The growth rate was normalized to the maximum values obtained across the full set of conditions. (C) Oligomer profiles and single-cell growth rates for simulated chambers after 20 h. Two scenarios are shown, taken from simulations in Figure 5B (I, *ρ* = 0.3 μm^-2^ and *K*_enz_ = 0.7 hr^-1^; II, *ρ* = 0.3 μm^-2^ and *K*_enz_ = 0.1 hr^-1^). The values of growth rates and oligomer concentrations are normalized to the maximum values observed in the simulations to allow comparison. (D) Growth rates and depolymerization rate of polysaccharides as a function of the diffusion coefficient of oligomers, polysaccharide, and enzymes. The diffusion coefficient was normalized by the diffusion coefficient in free water. Simulation parameters are described in detail in Table S1.

We first asked whether our model predicts similar density-dependent growth dynamics to those observed in the experimental observations. We simulated bacterial growth in the 2D chambers and calculated the mean growth rates over a wide range of initial cell densities and enzymatic activities. Consistent with our experimental observations, the model predicted that the growth rate of cells with lower enzymatic activity is higher at an increased cell density (Figure 5B). In comparison, cells with a higher enzymatic activity through increased secretions have higher growth rates at low densities (Figure 5B). To understand the biophysical mechanism underlying these effects, we examined the spatial profile of oligomer concentrations simulated by the model and found that lower enzymatic activity leads to the accumulation of oligomers and supports higher growth rates for cells regardless of their spatial arrangement. However, in higher enzymatic activity, oligomers arising as result of polysaccharide degradation are increasingly lost to diffusion without being available for bacterial cells to uptake (Figure 5C). Our results indicate that the rate of oligomer loss to diffusion and uptake plays a dominant role in shaping the nature of intercellular interactions in polysaccharide-degrading populations. We thus performed simulations where the diffusion coefficients of polymers and oligomers were systematically varied relative to diffusion in free water to yield low rates of diffusion such as in biofilm or cell aggregations and high rates of diffusion similar to boundary conditions in a quiescent region in contact with flow. The model suggests that a scenario with a diffusion coefficient that is an order of magnitude lower than that in bulk water results in the highest average growth rate for the bacterial population (Figure 5DE). A higher diffusion rate that resembles flow, while increasing the polymer degradation rate (Figure 5D), also enhances the loss of oligomers through diffusion, making them unavailable to bacterial cells and thus reducing the growth rate. A similar effect emerges at higher cell densities, where faster polymer degradation by cells at the opening of the growth chambers can result in loss of oligomers, thereby making them unavailable to other cells and leads to negative density dependent growth effects (figure 5C and E).

Therefore, our results indicate that enzymatic secretion capabilities can influence growth dynamics and intercellular interactions within bacterial populations growing on polysaccharides. Differences in enzyme secretion levels can arise as a result of strains harboring functionally distinct alginate lyases or as a result of distinct localization of alginate lyases^13,17–19^. For instance, *V. splendidus* 12B01 has alginate lyases that are largely intra-cellular or membrane-bound whereas alginate lyases of *V. splendidus* 13B01 are largely secreted into the extracellular environment^13,20^. In addition, strains can harbor distinct classes of alginate lyases and the breakdown products generated by these enzymes can differ^21^. Our findings suggest that cells that are genetically predisposed to produce low levels of enzymes or have predominantly membranebound alginate lyases can overcome growth disadvantages that exist in well-mixed environments by aggregating. These collective behaviors allow cells to generate 5-10-fold higher local cell densities enabling growth at maximal rates.

Polysaccharides like alginate or chitin exist predominantly as particulates in natural ecosystems^22^ and offer surfaces that enable bacterial cells to form dense colonies. In turn, aggregation behaviors increase local cell densities and enable intercellular interactions that can allow individuals to benefit from each other’s metabolic activities^11,23^. In contrast, strains that have increased enzymatic secretions can breakdown polysaccharides by adopting dispersed growth modes, that can facilitate migration of cells to new polysaccharide particles in nature. Additionally, in communities where multiple strains coexist^4,5,15,24^, the degradative activity of high enzyme secretors can create public goods that can positively influence the growth of not only the low enzyme producers but also strains that lack the ability to degrade polysaccharides and rely on cross-feeding of monomers from the polymer degraders^13,25,26^.

The cellular processes that link aggregative behaviors and polymer metabolism in marine bacterial populations are yet unclear. The formation of bacterial aggregates formation and production of polysaccharide-degrading enzymes both require substantial cellular resources and thus likely impose substantial metabolic costs on cells^7,27^. This will result in cells facing trade-offs between allocating resources to enzyme production versus aggregation, both of which enable cells to grow on polymers. A potential consequence is that marine polysaccharide degraders are constrained by trade-offs such that they forego enzymatic capabilities in favor of aggregation behaviors or alternately acquire genes encoding enzymes that can either be produced in large amounts and also be secreted instead of aggregative capabilities. While aggregation behaviors of bacterial cells are known to be coordinated through quorum sensing ^28,29^, the molecular mechanisms that regulate collective behaviors in response to the nutrient composition in the natural environment are yet unclear. Our work is a first step in this direction and provides a direct link between the dynamics of enzyme secretion, growth physiology and collective behaviors of polysaccharide degrading bacterial populations.

## Materials and Methods

### Bacterial strains, media and batch culture growth assays

Experiments were performed using *Vibrio splendidus* strains 13B01, 12B01, FF6 and 1S124; *Vibrio cyclitrophicus* strains 1F175, 1F111, ZF270 and ZF28; and *Vibrio sp.* F13 strains 9CS106, 9ZC13, 9ZC77 and ZF57. The isolation conditions and the genes encoding alginate lyases in these strains have been described previously^13^. Prior to experiments, strains were cultured in Marine Broth (MB medium, DIFCO) and grown for 18 h at 25 °C. Cells from these cultures were then used for batch culture growth kinetic assays in Tibbles–Rawling Minimal Medium^13,30^ (TR medium) containing 0.1% (weight/volume) alginate (Sigma Aldrich). Sterile alginate solution was prepared using nanopure water and filter sterilized using 0.40 μm surfactant-free cellulose acetate filters (Corning). For growth assays with enzyme supplementation, 0.05 units ml^-1^ of alginate lyase (Sigma) was added to TR medium containing 0.1% alginate. Batch growth kinetic assays were performed in 96-well plates and growth was measured using a micro-well plate reader (Biotek). For this, 1 ml of cell suspension from a culture growing in MB medium was centrifuged (13000 rpm for 2 min) in a 2 ml microfuge tube. The supernatant was discarded and the cell pellet was washed twice with TR medium without any carbon source. The cell pellet was resuspended in 500 μl of TR medium without carbon source and the optical density measured and adjusted to 0.1 OD. For the experiments, 5 μl of this OD-adjusted cell suspension was used to seed wells containing 195 μl TR minimal medium with the carbon source. Cells were allowed to grow for 40 h at 25 °C and OD measurements taken every 30 minutes. Each experiment was replicated 3–6 times.

### Enzyme secretion assay

An agarose plate assay^16^ was used to test the ability of strains to secrete alginate lyases. Prior to experiments, cultures were grown for 18 h in MB medium and 1 ml of cell suspension was centrifuged (13000 rpm for 2 min) in a 2 ml microfuge tube. The supernatant was discarded and the cell pellet was washed twice with TR medium without any carbon source and then centrifuged. The cell pellet was suspended in 1 ml of TR medium without carbon source and the optical density measured and adjusted to 0.1 OD. For the assays, 50 μl of this culture was spotted on plates that were made using TR medium containing 0.1% (w/v) alginate and 1% agarose (Applichem). Colonies were allowed to grow for 30 h at 25 °C and then the plates were flooded with 2% Gram’s iodine. Excess iodine was discarded and the plate imaged using a 12MP iPhone camera. If cells secreted alginate lyases, then a distinct clearance zone was formed, the diameter of which was measured using a ruler.

### Microfluidics and time-lapse microscopy

The setup of microfluidics experiments has been described in detail previously^11,23^. Cell growth and behavior was imaged within chambers measuring 60–120 × 60 × 0.56 μm (length × width × height). In these chambers, cells could adhere to the glass surface and experienced TR medium with 0.1% alginate that diffused into the lateral flow channels. Microscopy imaging was performed using an IX83 inverted microscope system with automated stage controller (Marzhauser Wetzlar), shutter, and laser-based autofocus system (Olympus ZDC 2). Chambers were imaged in parallel on the same PDMS chip, capturing phase-contrast images of each position every 8–10 min. Images were acquired using an UPLFLN 100× oil immersion objective (Olympus) and an ORCA-flash 4.0 v2 or v4 sCMOS cameras (Hamamatsu, Japan). For image acquisition, the CellSens 1.18 and higher software package (Olympus) were used. The microscopy units and PDMS chip were maintained at 25 °C using a cellVivo microscope incubation system (Pecon GmbH).

### Image analysis

Cell segmentation and tracking was performed in ilastik (v1.2 and newer)^31^. The phase-contrast images were aligned using SuperSegger^32^. Images were cropped at the boundaries of each microfluidic chamber. Growth properties (cell area over time) and spatial locations (*x* and *y* coordinates) were directly derived using the ilastik Tracking plugin and using a custom algorithm (available in repository).

### Datasets and statistical analysis

Growth curves were analyzed in Python (v3.8) using the AMiGA software^33^ and GraphPad Prism v9 (GraphPad Software). The microscopy dataset set consists of seven chambers for each strain. These are grouped into two biological replicates wherein each biological replicate is fed by medium through one unique channel in a microfluidic chip, with three chambers fed by one channel and four chambers by another. Cells with negative growth rates (13B01 = 20%, 12B01 = 33%, 1F111 = 36%, ZF270 = 24%, 9ZC13 = 26%, 9ZC77 = 27%) were excluded from the analysis after visual curation, and represent artefacts, mistakes in linking during the segmentation process or non-growing deformed cells. In total, 2524 cells were analyzed for 13B01, 11547 cells for 12B01, 3227 cells for 1F111, 24444 cells for ZF270, 8910 cells for 9ZC13 and 24335 cells for 9ZC77. Each chamber was treated as an independent replicate. Each figure depicts medians of the seven chambers for each strain. Detailed equations for non-linear regression models for growth rate as a function of cell density are provided in Supplementary Methods. Comparisons were considered statistically significant when *P* > 0.05. FDR corrections were applied when multiple *t* tests were performed for the same dataset. Measures of effect size are represented by the *R*^2^ or eta^2^ value. All statistical analyses were performed in either GraphPad Prism v 9.0 (GraphPad Software) or Rstudio v1.1.463 (Rstudio inc). All data analysis scripts are deposited along with the raw data.

## Data Availability

All curated image analysis datasets and source data for figures will be deposited in Zenodo data repository upon publication.

## Supplemental Figure Legends

**Figure S1: Strains display distinct growth dynamics on the marine polysaccharide alginate.** Populations of strains belonging to (**A**) *Vibrio splendidus*, (**B**) *Vibrio cyclitrophicus* and (**C**) *Vibrio sp.* F13 were grown in the same concentration (%weight/volume) of the polysaccharide alginate and population size (optical density at 600 nm) were measured every 15 min over 36 h. Colours represent different strains within each species. Circles and error bars indicate the mean of the individual measurements for each population (*n*_populations_ = 3) and the 95% confidence interval (CI), respectively.

**Figure S2: Alginate lyase assay on alginate agar plates.** (**A**) *Vibrionaceae* strains were screened for alginate lyase secretion by allowing them to grow for 36 h on alginate agar plates, then stained using iodine to measure the diameter of the colony spot (black circle) and the diameter of the halo (white circle) produced by alginate digestion (see methods). Shown are representative images of plates with halos after staining. (**B**) Strains vary in their secreted alginate lyase production when growing on alginate plates. The diameter of the halo measured using an iodine assay after a 36-hour growth cycle is used as a proxy for alginate lyase production and activity (see Materials and Methods, Figure S2). The bars represent the mean of replicates of each ecotype (*n* = 3) while error bars indicate the 95% confidence intervals (CI). Letters indicate statistically significant differences between strains within each species (Kruskal–Wallis (KW) test and Dunn’s post-hoc test; *V. splendidus: P* = 0.0001, KW statistic = 21.04; *V. cyclitrophicus: P* = 0.0002, KW statistic = 19.95, *V. sp* F13: *P* < 0.0001, KW statistic = 23.00).

**Figure S3: Growth properties of *Vibrionaceae* strains on alginate and alginate supplemented with alginate lyases.** (**A**) Maximum optical density (OD600) over a growth cycle of ecotype populations belonging to *Vibrio splendidus*, *Vibrio cyclitrophicus* and *Vibrio sp.* F13. Cells were grown in the same concentration (%weight/volume) of the polysaccharide alginate alone (circles) or supplemented with alginate lyases (triangles). **(B)** Time to reach exponential growth phase when grown on alginate (circles) or alginate supplemented with external alginate lyases (triangles). (**C**) Correlation between the reduction in time to exponential growth when supplemented with alginate lyases and the halo diameter when grown on an alginate agar plate across the 12 strains. The line indicates the fit of a linear regression model (slope *R^2^* = 0.65, *R* = 0.58, *P* < 0.0001; Spearman’s correlation *r* = 0.61, *P* = 0.03). Circles and triangles (black, *Vibrio splendidus;* green, *Vibrio cyclitrophicus;* and purple, *Vibrio sp.* F13) represent the mean of the populations (*n*_populations_ = 3), error bars indicate 95% confidence intervals.

**Figure S4: Aggregation dynamics of *Vibrionaceae* strains on alginate in microfluidic growth chambers.** Strains that are low secretors (blue boxes) of alginate lyases reach a higher density within the microfluidic growth chambers compared to strains that secrete higher levels (orange boxes) of alginate lyases. Asterisks indicate statistically significant differences between the two strains within a species (unpaired *t*-tests, *n*_chambers_ = 7 in each case; *V. splendidus, P* < 0.0001, *t* = 17.30, df = 12, *R*^2^ (eta squared) = 0.96; *V. cyclitrophicus, P* < 0.0001, *t* = 10.66, df = 12, *R*^2^ (eta squared) = 0.90; *V. sp* F13, *P* < 0.0001, *t* = 14.63, df = 12, *R*^2^ (eta squared) = 0.94). Box plots extend from the 25th to 75th percentiles and whiskers indicate the 10th and 90th percentiles.

**Figure S5: Distribution of single cell growth rates of *Vibrionaceae* strains.** High enzyme secreting strains tend to reach maximum growth rates earlier than low alginate lyase secreting strains while growing on alginate. Smoothed single cell growth rates of (A) *V. splendidus,* (B) *V. cyclitrophicus* and (C) *V. sp.* F13 cells across different time bins. Cells were binned into 2 h intervals based on their birth times (bins: 0–2.66 h, 2.67–5.32 h, 5.32–7.98 h, 7.99–10.64 h, 10.65–13.30 h, 13.31–15.96 h, 15.97–18.62 h, 18.63–21.28 h, 21.29–23.94 h and 23.95–26.60 h) and the distribution plotted as ridges. Orange distributions indicate the high secretors of alginate lyase (A, 13B01; B,1F111; and C, 9ZC13) while the blue distributions indicate the relatively low secretors of alginate lyases (A, 12B01; B, ZF270; and C, 9ZC77). White vertical lines indicate the median of each distribution.

**Figure S6: Growth dynamics of strains in well-mixed and microfluidic growth environments** (**A**) Strains within each species differ in the median growth rate of cells present during the final birth time bin (23.95–26.60 h) within chambers (unpaired *t*-tests, *n*_chambers_ = 7 in each case; *V. splendidus, P* < 0.05, *t* = 2.27, df = 10, *R*^2^ (eta squared) = 0.34; *V. cyclitrophicus, P* < 0.0001, *t* = 4.5, df = 12, *R*^2^ (eta squared) = 0.63; *V. sp* F13, *P* < 0.001, *t* = 3, df = 12, *R*^2^ (eta squared) = 0.45). (**B**) Growth rate of the low enzyme secreting ecotype relative to the growth rate of the high enzyme secreting ecotype within each species. *S* and *M* indicate growth rates measured in shaken well-mixed and microfluidic environments, respectively. Asterisks indicate statistically significant differences between growth environments within a species (Welch’s *t*-test, *n*_popuiations_ = 3 (for well-mixed batch assays), *n*_chambers_ = 7 (for microfluidics assays); *V. splendidus, P* < 0.001, *t* = 22.90, df = 5.33, *R*^2^ (eta squared) = 0.98; *V. cyclitrophicus, P*> 0.05, *t* = 1.1, df = 6.07, *R*^2^ (eta squared) = 0.16; *V. sp* F13, *P* < 0.001, *t* = 5.60, df = 6.26, *R*^2^ (eta squared) = 0.83).

**Figure S7: Inoculum cell density influences the growth dynamics of *Vibrionaceae* strains.** Maximum growth rate (h^-1^) of populations of (A) *V. splendidus,* (B) *V. cyclitrophicus* and (C) *V. sp.* F13 as a function of initial cell density (cfu ml^-1^). Circles indicate the mean measurements for each biological replicate (*n* = 3) while error bars indicate the confidence intervals.

## Supplemental Table Legend

Table S1. Physiological and chemical parameters for microbial growth, metabolism and nutrient concentrations in the individual-based model.

**Supplementary videos:**

**Supplemental Video 1 (separate file).** Time-lapse of *Vibrio splendidus* 13B01 cells within a representative microfluidics chamber fed with 0.1% alginate as the sole carbon source. Images were captured every 8 minutes.

**Supplemental Video 2 (separate file).** Time-lapse of *Vibrio splendidus* 12B01 cells within a representative microfluidics chamber fed with 0.1% alginate as the sole carbon source. Images were captured every 8 minutes.

**Supplemental Video 3 (separate file).** Time-lapse of *Vibrio cyclitrophicus* 1F111 cells within a representative microfluidics chamber fed with 0.1 % alginate as the sole carbon source. Images were captured every 8 minutes.

**Supplemental Video 4 (separate file).** Time-lapse of *Vibrio cyclitrophicus* ZF270 cells within a representative microfluidics chamber fed with 0.1 % alginate as the sole carbon source. Images were captured every 8 minutes.

**Supplemental Video 5 (separate file).** Time-lapse of *Vibrio sp. F13* 9ZC13 cells within a representative microfluidics chamber fed with 0.1% alginate as the sole carbon source. Images were captured every 8 minutes.

**Supplemental Video 6 (separate file).** Time-lapse of *Vibrio sp. F13* 9ZC77 cells within a representative microfluidics chamber fed with 0.1% alginate as the sole carbon source. Images were captured every 8 minutes.

## Acknowledgements

We thank Daan Kiviet and Susan Schlegel for designing the microfluidic growth chambers, and past and present members of the Microbial Systems Ecology group for feedback. This research was funded by an ETH fellowship and a Marie Curie Actions for People COFUND program fellowship (FEL-37-16-1) to GD; a Career Seed Grant from ETH Zurich to GD; the Simons Foundation Collaboration on Principles of Microbial Ecosystems (PriME #542389 and #542395) to MA, OC and RS; a grant from the Swiss National Science Foundation (31003A_169978) to MA; and by ETH Zurich and Eawag.

## Author Contributions

GD conceived the research along with MA, AE and OC. AE developed the mathematical model and simulations along with OC. GD designed and performed all experiments. AS developed the algorithm to compute growth rates from the cell segmentation pipeline. GD analyzed the data with inputs from AE, JK, OC, RS and MA. MD developed the alginate lyase detection assay based on previous literature. GD and AE wrote the manuscript with inputs from JK, AS, MD, RS, OC and MA.

