## Supplementary information file for "Intercellular collectivity is governed by enzyme secretion strategies in marine polysaccharide degrading bacteria"

**This file includes:**

Supplementary Methods

Supplementary Figures 1 to 7

Supplementary Videos 1 to 6

Supplementary Text

Supplementary Table 1

Supplementary References

**Supplementary Figures**:

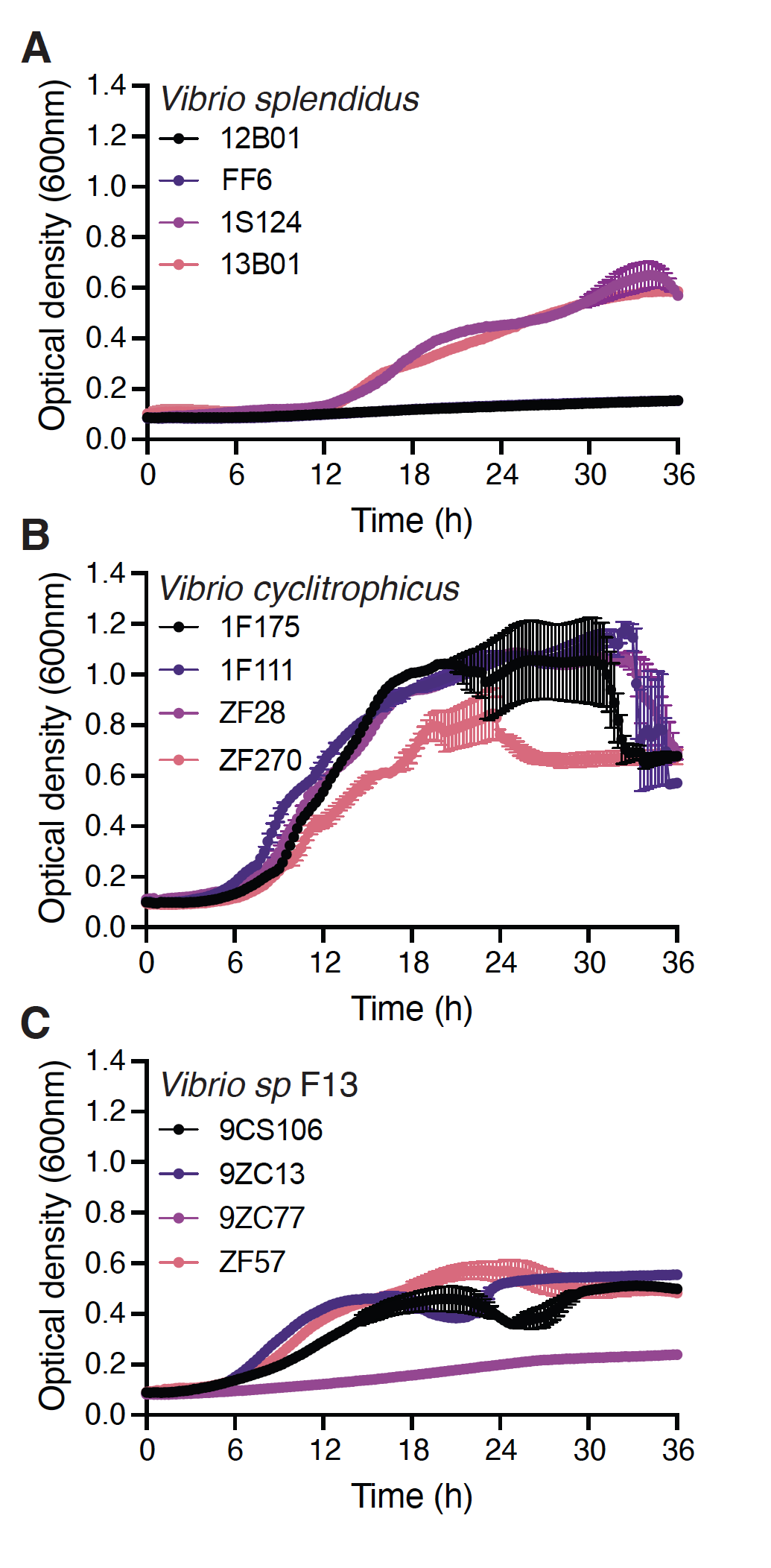

**Figure S1: Strains display distinct growth dynamics on the marine polysaccharide alginate.** Populations of strains belonging to (**A**) *Vibrio splendidus*, (**B**) *Vibrio cyclitrophicus* and (**C**) *Vibrio sp.* F13 were grown in the same concentration (%weight/volume) of the polysaccharide alginate and population size (optical density at 600 nm) were measured every 15 min over 36 h. Colours represent different strains within each species. Circles and error bars indicate the mean of the individual measurements for each population (*n*_populations_ = 3) and the 95% confidence interval (CI), respectively.

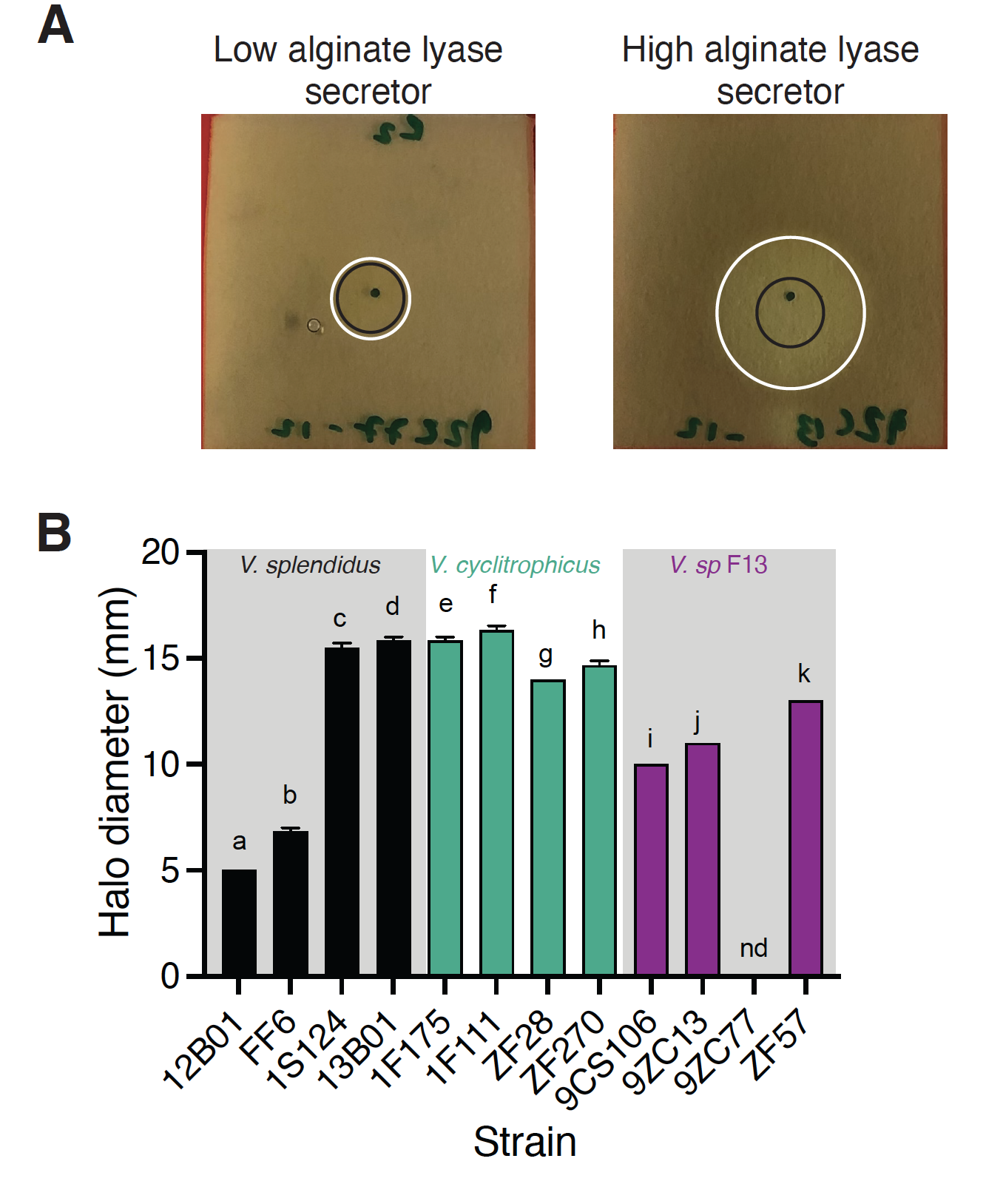

**Figure S2: Alginate lyase assay on alginate agar plates.** (**A**)*Vibrionaceae* strains were screened for alginate lyase secretion by allowing them to grow for 36 h on alginate agar plates, then stained using iodine to measure the diameter of the colony spot (black circle) and the diameter of the halo (white circle) produced by alginate digestion (see methods). Shown are representative images of plates with halos after staining. (**B**) Strains vary in their secreted alginate lyase production when growing on alginate plates. The diameter of the halo measured using an iodine assay after a 36-hour growth cycle is used as a proxy for alginate lyase production and activity (see Materials and Methods, Figure S2). The bars represent the mean of replicates of each ecotype (*n* = 3) while error bars indicate the 95% confidence intervals (CI). Letters indicate statistically significant differences between strains within each species (Kruskal–Wallis (KW) test and Dunn’s post-hoc test; *V. splendidus*: *P* = 0.0001, KW statistic = 21.04; *V. cyclitrophicus*: *P* = 0.0002, KW statistic = 19.95, *V. sp* F13: *P* < 0.0001, KW statistic = 23.00)

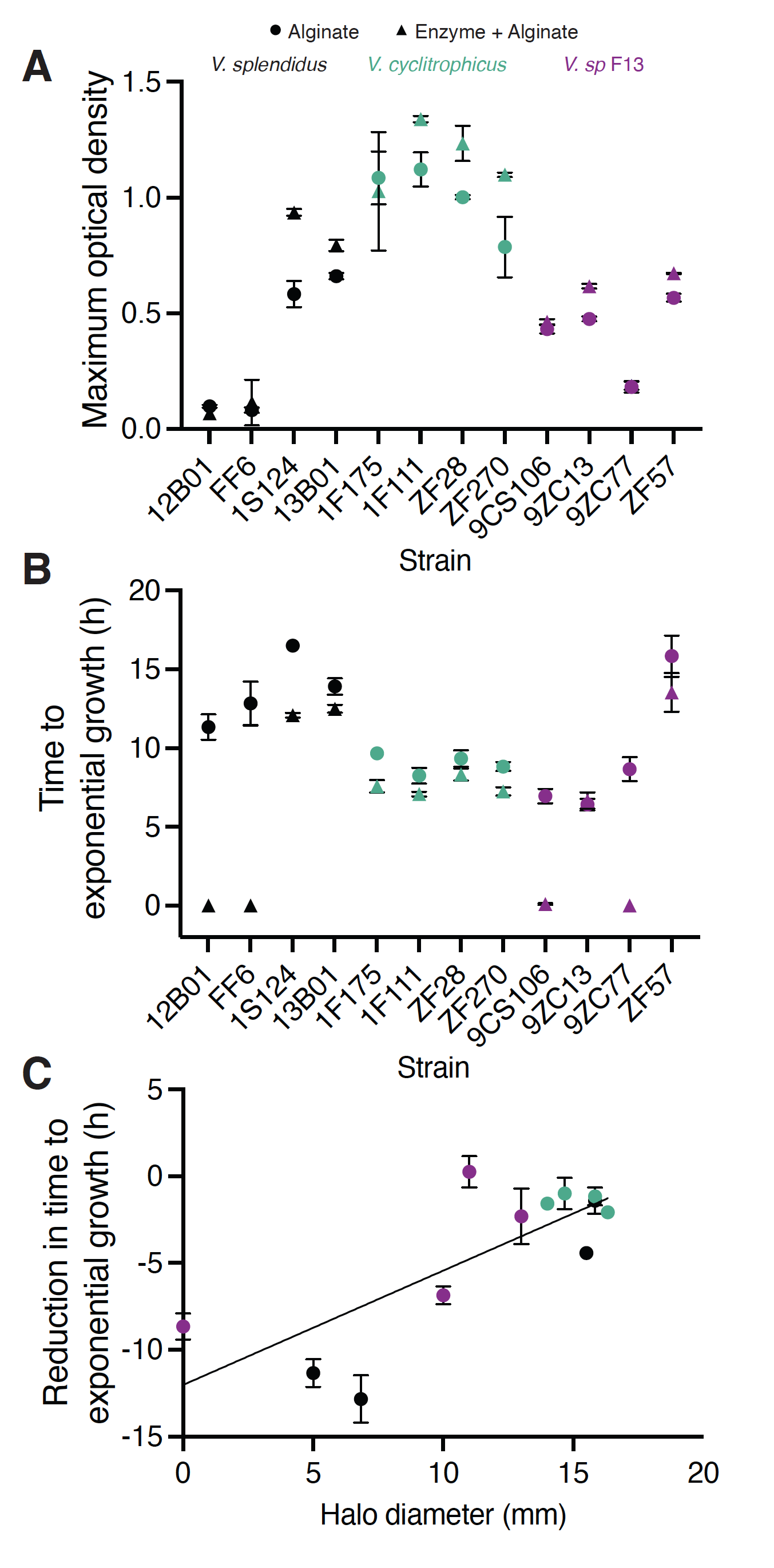

**Figure S3: Growth properties of *Vibrionaceae* strains on alginate and alginate supplemented with alginate lyases.** (**A**) Maximum optical density (OD600) over a growth cycle of ecotype populations belonging to *Vibrio splendidus*, *Vibrio cyclitrophicus* and *Vibrio sp.* F13. Cells were grown in the same concentration (%weight/volume) of the polysaccharide alginate alone (circles) or supplemented with alginate lyases (triangles). **(B)** Time to reach exponential growth phase when grown on alginate (circles) or alginate supplemented with external alginate lyases (triangles). (**C**) Correlation between the reduction in time to exponential growth when supplemented with alginate lyases and the halo diameter when grown on an alginate agar plate across the 12 strains. The line indicates the fit of a linear regression model (slope = 0.65, *R*^2^ = 0.58, *P* < 0.0001; Spearman’s correlation *r* = 0.61, *P* = 0.03). Circles and triangles (black, *Vibrio splendidus;* green, *Vibrio cyclitrophicus;* and purple, *Vibrio sp.* F13) represent the mean of the populations (*n*_populations_ = 3), error bars indicate 95% confidence intervals.

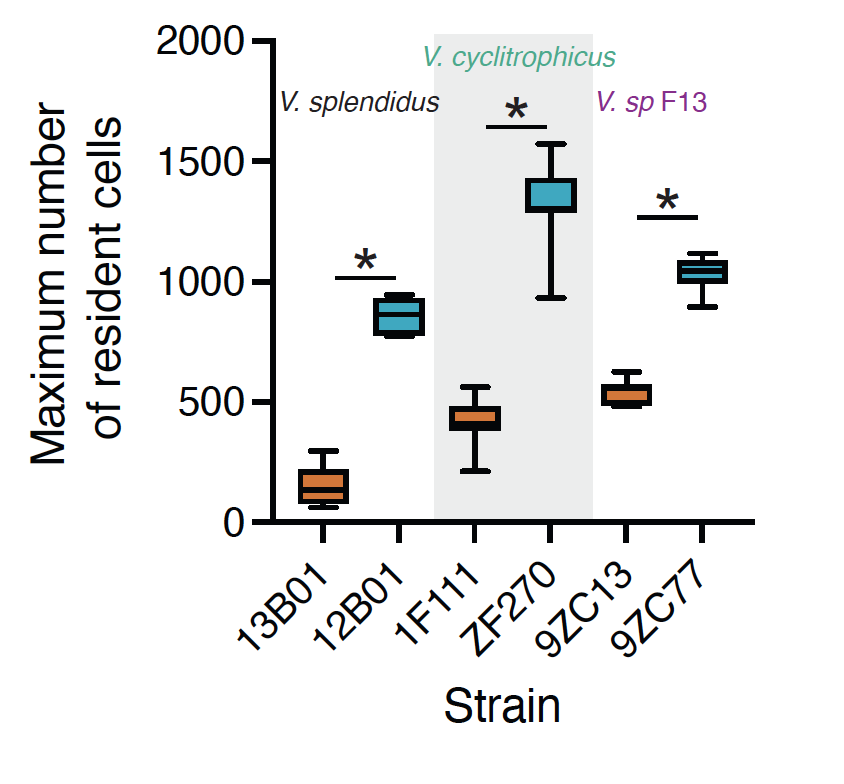

**Figure S4: Aggregation dynamics of *Vibrionaceae* strains on alginate in microfluidic growth chambers.** Strains that are low secretors (blue boxes) of alginate lyases reach a higher density within the microfluidic growth chambers compared to strains that secrete higher levels (orange boxes) of alginate lyases. Asterisks indicate statistically significant differences between the two strains within a species (unpaired *t*-tests, *n*_chambers_ = 7 in each case; *V. splendidus*, *P* < 0.0001, *t* = 17.30, df = 12, *R*^2^ (eta squared) = 0.96; *V. cyclitrophicus*, *P* < 0.0001, *t* = 10.66, df = 12, *R*^2^ (eta squared) = 0.90; *V. sp* F13, *P* < 0.0001, *t* = 14.63, df = 12, *R*^2^ (eta squared) = 0.94). Box plots extend from the 25th to 75th percentiles and whiskers indicate the 10th and 90th percentiles.

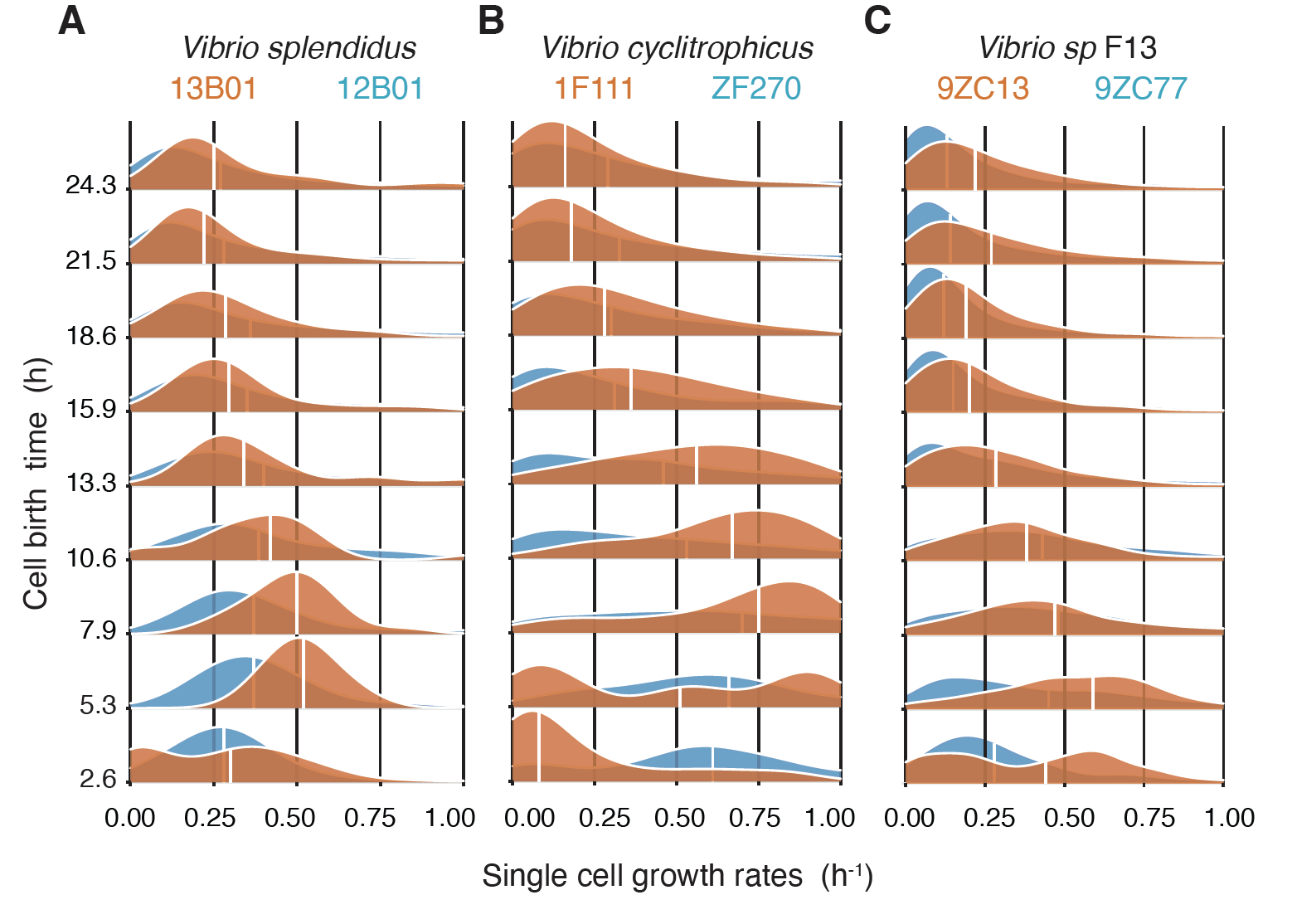

**Figure S5: Distribution of single cell growth rates of *Vibrionaceae* strains.** High enzyme secreting strains tend to reach maximum growth rates earlier than low alginate lyase secreting strains while growing on alginate. Smoothed single cell growth rates of (A) *V. splendidus*, (B) *V. cyclitrophicus* and (C) *V. sp.* F13 cells across different time bins. Cells were binned into 2 h intervals based on their birth times (bins: 0–2.66 h, 2.67–5.32 h, 5.32–7.98 h, 7.99–10.64 h, 10.65–13.30 h, 13.31–15.96 h, 15.97–18.62 h, 18.63–21.28 h, 21.29–23.94 h and 23.95–26.60 h) and the distribution plotted as ridges. Orange distributions indicate the high secretors of alginate lyase (A, 13B01; B,1F111; and C, 9ZC13) while the blue distributions indicate the relatively low secretors of alginate lyases (A, 12B01; B, ZF270; and C, 9ZC77). White vertical lines indicate the median of each distribution.

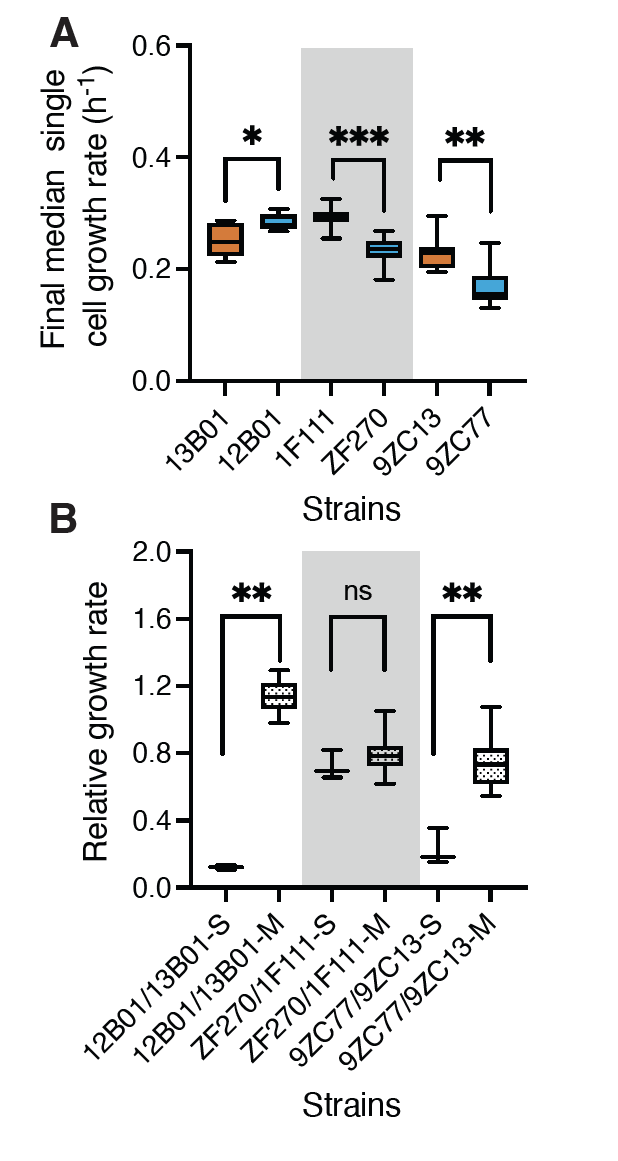

**Figure S6: Growth dynamics of strains in well-mixed and microfluidic growth environments** (**A**) Strains within each species differ in the median growth rate of cells present during the final birth time bin (23.95–26.60 h) within chambers (unpaired *t*-tests, *n*_chambers_ = 7 in each case; *V. splendidus,* *P*< 0.05, *t* = 2.27, df = 10, *R*^2^ (eta squared) = 0.34; *V. cyclitrophicus,* *P* < 0.0001, *t* = 4.5, df = 12, *R*^2^ (eta squared) = 0.63; *V. sp* F13, *P* < 0.001, *t* = 3, df = 12, *R*^2^ (eta squared) = 0.45). (**B**) Growth rate of the low enzyme secreting ecotype relative to the growth rate of the high enzyme secreting ecotype within each species. *S* and *M* indicate growth rates measured in shaken well-mixed and microfluidic environments, respectively. Asterisks indicate statistically significant differences between growth environments within a species (Welch’s *t*-test, *n*_populations_ = 3 (for well-mixed batch assays), *n*_chambers_ = 7 (for microfluidics assays); *V. splendidus,* *P* < 0.001, *t* = 22.90, df = 5.33, *R*^2^ (eta squared) = 0.98; *V. cyclitrophicus,* *P* > 0.05, *t* = 1.1, df = 6.07, *R*^2^ (eta squared) = 0.16; *V. sp* F13, *P* < 0.001, *t* = 5.60, df = 6.26, *R*^2^ (eta squared) = 0.83).

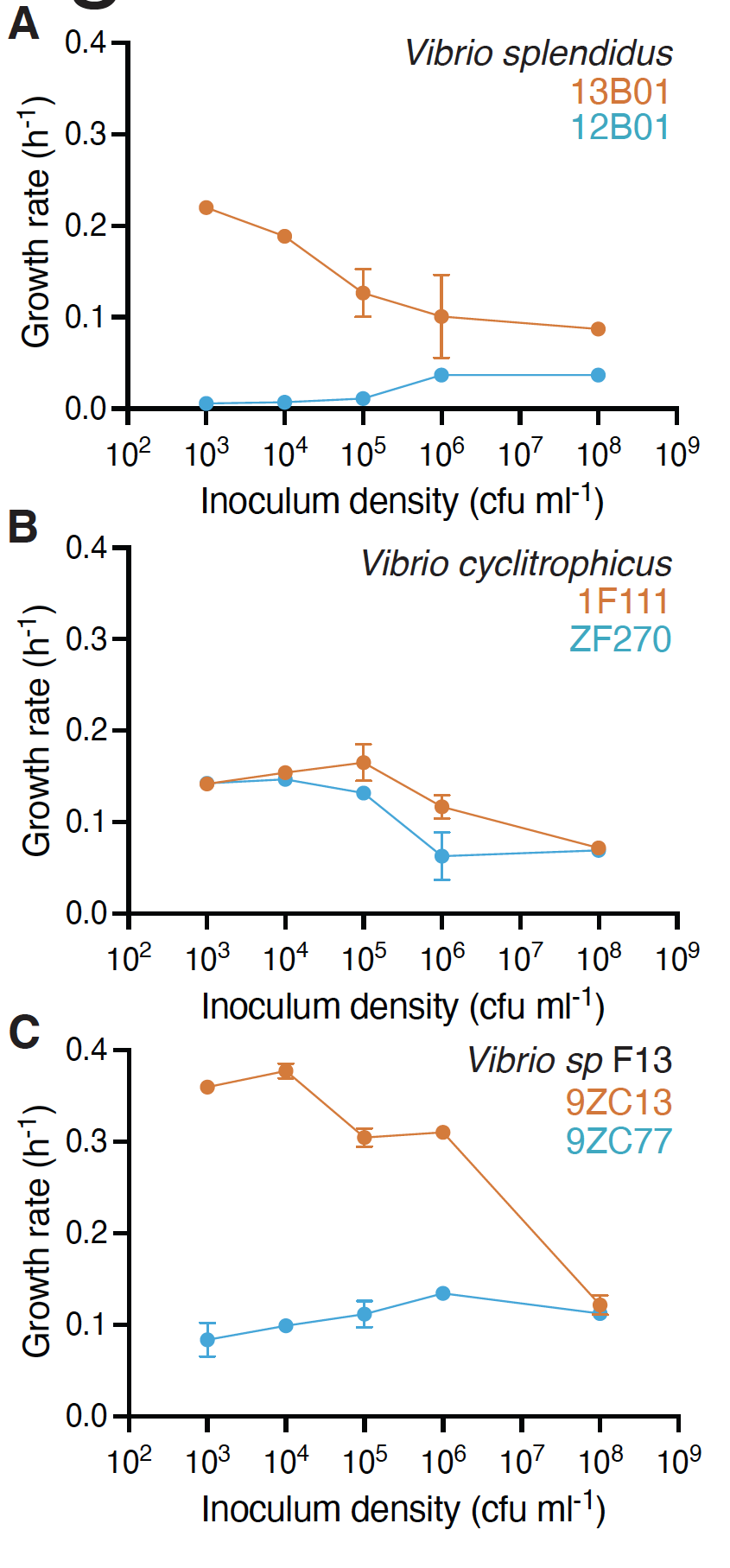

**Figure S7: Inoculum cell density influences the growth dynamics of *Vibrionaceae* strains.** Maximum growth rate (h^-1^) of populations of (A) *V. splendidus*, (B) *V. cyclitrophicus* and (C) *V. sp.* F13 as a function of initial cell density (cfu ml^-1^). Circles indicate the mean measurements for each biological replicate (*n* = 3) while error bars indicate the confidence intervals.

**Supplementary videos:**

Cells from *Vibrionaceae* populations growing in microfluidic growth chambers. Microscopy images were taken using phase contrast every 8 minutes. Cells are false colored based on the identity of their progenitor cells. Frame rate for all movies is 10 images per second. Scale bar is 10 microns. Cells whose divisional history cannot be tracked are show in blue.

**Supplemental Video 1 (separate file).** Time-lapse of *Vibrio splendidus* 13B01 cells within a representative microfluidics chamber fed with 0.1% alginate as the sole carbon source. Images were captured every 8 minutes.

**Supplemental Video 2 (separate file).** Time-lapse of *Vibrio splendidus* 12B01 cells within a representative microfluidics chamber fed with 0.1% alginate as the sole carbon source. Images were captured every 8 minutes.

**Supplemental Video 3 (separate file).** Time-lapse of *Vibrio cyclitrophicus* 1F111 cells within a representative microfluidics chamber fed with 0.1% alginate as the sole carbon source. Images were captured every 8 minutes.

**Supplemental Video 4 (separate file).** Time-lapse of *Vibrio cyclitrophicus* ZF270 cells within a representative microfluidics chamber fed with 0.1% alginate as the sole carbon source. Images were captured every 8 minutes.

**Supplemental Video 5 (separate file).** Time-lapse of *Vibrio sp. F13* 9ZC13 cells within a representative microfluidics chamber fed with 0.1% alginate as the sole carbon source. Images were captured every 8 minutes.

**Supplemental Video 6 (separate file).** Time-lapse of *Vibrio sp. F13* 9ZC77 cells within a representative microfluidics chamber fed with 0.1% alginate as the sole carbon source. Images were captured every 8 minutes.

**Supplementary text:**

**Mathematical modelling of bacterial growth in microfluidic growth chambers**

The following represents derivation of mathematical formulations and procedures to model microbial growth, enzymatic activity, and diffusion processes in 2D chamber system.

**S1.1 Growth and division**

In this model, the growth kinetics of individual cells,
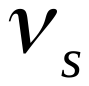
are dictated by Monod-type kinetics given as

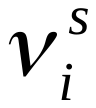
=
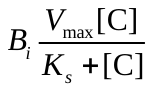
 (S1)

where
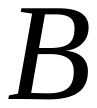
is the cell dry mass of an individual cell, *i* and
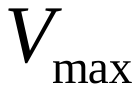
is the maximum carbon uptake and defined as:
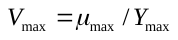
 (maximum specific growth rate / carbon conversion yield to biomass). *Ks* is the half-saturation constant for oligomer. [C] is oligomer concentration and is assumed as the primary limiting substrate for bacterial growth. We kept all other nutrients (e.g., oxygen and nitrogen) available at sufficient levels for bacterial growth.

The actual biomass accumulation (
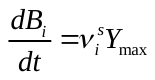
) and maintenance of an individual cell (
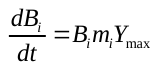
) are linearly correlated with cell dry mass and therefore the new growth rate of individual cells (
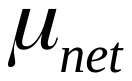
) is given as:

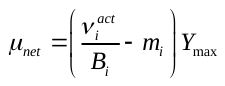
 (S2)

In the individual based model, each cell may double to two daughter cells when a certain amount of carbon has been taken up^1,2^. The minimum volume of the individual cell at the threshold for the division (
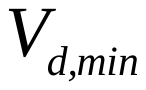
) is given from the descriptive Donachie model ^1,2^

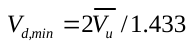
 (S3)

where
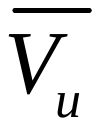
 is the median volume of the individual cell. The cell will divide into two identical (half-volume) cells if reaches
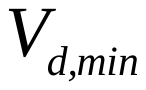
. In the simulation, cylindrical cells are assumed. Under limited nutrient conditions, the actual growth rate of an individual cell is restricted by available amount of oligomer within the domain. In each time step, the total amount of oligomer at the given grid is calculated and compared with the amount of oligomer needed for bacterial growth that is calculated from Monod kinetics. If the predicted growth rate from the Monod kinetics requires more oligomer than the available amount, the growth rate is adjusted to satisfy mass conservation. The parameters of the growth kinetics used for the simulations are provided in Table SX.

**S1.2 Enzymatic activity**

Extracellular enzymes at individual cell level are modeled by assuming that the enzyme is broadcasted and diffused to the extracellular environment around the individual cells. The enzyme secretion rate (
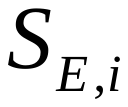
) is assumed to be linearly correlated with cell biomass,
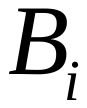
 of a bacterial cell, *i*:

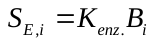
 (S4)

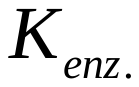
is the enzyme production rate.

1.1 **Oligomer and polymer diffusions along aggregate radius**

The model explicitly simulates the diffusion of oligomers and polymers within the 2D chamber. For oligomers, an absorbing condition (zero concentration) is assumed at periphery of the chamber that simulates loss of oligomers to constant flow rate. Diffusion of oligomers is modelled based on Fick’s law of diffusion. Reaction-diffusion equation is then numerically solved by finite-difference method:

$\frac{d[C]\left( x,y \right)}{dt}=K_{p}.\left[ E \right] \left( x,y \right).\left[ P \right]\left( x,y \right)-B.V_{max}.\frac{\left[ C \right]\left( x,y \right)}{\left[ C \right]\left( x,y \right)+K_{s}}+D_{[C]}\left( \frac{\partial^{2}[C]\left( x,y \right)}{{\partial x}^{2}}+\frac{\partial^{2}[C]\left( x,y \right)}{{\partial y}^{2}} \right)$	(S5)

where [*E*] is the concentration of enzyme. *K_p_*_._ is polymer lability that defines how many grams of oligomers are released per gram of enzyme acting on the polymer surface per unit of time. *B* is the total biomass ^3^. *D*_[_*_C_*_]_ is the diffusion coefficient of oligomers.

Similar to oligomers, polymer diffusion is given as:

$\frac{d[P]\left( x,y \right)}{dt}=-K_{p}.\left[ E \right] \left( x,y \right).\left[ P \right]\left( x,y \right)+D_{[P]}\left( \frac{\partial^{2}[P]\left( x,y \right)}{{\partial x}^{2}}+\frac{\partial^{2}[P]\left( x,y \right)}{{\partial y}^{2}} \right)$	(S6)

The physiological and chemical parameters used in the simulations are represented in Table S1.

**Supplementary table:**

**Table S1. Physiological and chemical parameters for microbial growth, metabolism and nutrient concentrations in the individual-based model.**

| *Parameters* | *Values (Units)* |
| --- | --- |
| 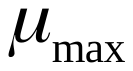: *maximum growth rate* (*hr^-1^*) | 0.1^*^ |
| 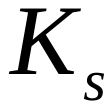: *half saturation (mg/L)* | 0.1^*^ |
| 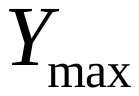 : *growth yield* (*gr dry mass/gr substrate*) | 0.5^*^ |
| *cell maintenance* | 0.1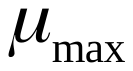^ſ^ |
| *cell size* (*µm*) | 1 ^ſ^ |
| *ρ: cell density* (*mg L^-1^*) | 2.9×10^5 ſ^ |
| *V_1D_ : cell velocity at bulk solution (µm/s)* | 0^*^ |
| *V_u_ : median cell volume (fl)* | 0.4 ^ſ^ |
| *K_p_ polymer lability (hr^-1^)* | 100^\|\|^ |
| 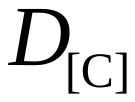 *diffusion coefficient of oligomers in bulk liquid (m^2^hr^-1^)* | 2.4×10^-6^ |
| *D_[p]_^.^ Polymer diffusion coefficient in bulk liquid (m^2^hr^-1^)* | 1.7×10^-5^ |

^ſ	1^

*^||^Enzyme Database – BRENDA: mean observed value for alginate*

**Model assumption*
